# Exploring the kinetic selectivity of drugs targeting the β_1_-adrenoceptor

**DOI:** 10.1101/2021.08.31.458064

**Authors:** David A. Sykes, Mireia Jiménez-Rosés, John Reilly, Robin A. Fairhurst, Steven J. Charlton, Dmitry B. Veprintsev

**Author notes:** **Corresponding author(s):** Dr David A. Sykes, School of Life Sciences, Queen’s Medical Centre, University of Nottingham, Nottingham NG7 2UH, Prof Steven Charlton, School of Life Sciences, Queen’s Medical Centre, University of Nottingham, Nottingham NG7 2UH, Prof Dmitry B. Veprintsev, School of Life Sciences, Queen’s Medical Centre, University of Nottingham, Nottingham NG7 2UH.

## Abstract

In this study, we report the β_1_-adrenoceptor binding kinetics of several clinically relevant β_1/2_-adrenoceptor (β_1/2_AR) agonists and antagonists. We demonstrate that the physicochemical properties of a molecule directly affect its kinetic association rate (*k*_on_) and affinity for the target. In contrast to our findings at the β_2_-adrenoceptor, a drug’s immobilized artificial membrane partition coefficient (K_IAM_), reflecting both hydrophobic and electrostatic interactions of the drug with the charged surface of biological membranes, was no better predictor than simple hydrophobicity measurements such as log P or logD_7.4_, characterized by a distribution between water and a non-aqueous organic phase (e.g. n-octanol) at predicting association rate. Overall, this suggests that hydrophobic interactions rather than a combination of polar and hydrophobic interactions play a more prominent role in dictating the binding of these ligands to the β_1_-adrenoceptor.

Using a combination of kinetic data, detailed structural and physicochemical information we rationalize the above findings and speculate that the association of positively charged ligands at the β_1_AR is curtailed somewhat by its predominantly neutral/positive charged extracellular surface. Consequently, hydrophobic interactions in the ligand binding pocket dominate the kinetics of ligand binding. In comparison at the β_2_AR, a combination of hydrophobicity and negative charge attracts basic, positively charged ligands to the receptor’s surface promoting the kinetics of ligand binding. Additionally, we reveal the potential role kinetics plays in the on-target and off-target pharmacology of clinically used β-blockers.

## Introduction

The β_1_-adrenergic receptor is involved in the sympathetic nervous systems control of the circulation mainly via its stimulatory signaling effects on the heart. So called β-blockers, also known as β-adrenergic receptor blockers, decrease heart rate and blood pressure through their actions at β_1_-adrenoceptors and are therefore useful in the treatment of hypertension and heart failure. The promiscuous binding of β-adrenergic ligands to the closely related subtypes of the β-adrenoceptor (e.g., the β_2_-adrenergic receptor) leads to commonly observed side effects such as fatigue, decreased peripheral circulation and increased airway resistance. This cross reactivity has long been considered a risk factor in asthmatic patients, with deaths attributed to β-blockers use in the very early years of their use, leading to the recommendation that non-selective β-blockers be avoided in asthmatic subjects (Harries, 1981; Harrington, 2020). The heart is not the only target for the therapeutic action of β-blockers, reflected in their use in the treatment of migraine, essential tremor, pheochromocytoma, thyrotoxicosis anxiety and the most common form of glaucoma (Baker et al., 2011; Harrington, 2020). More recently reports suggest a role for β-blockers in the treatment of certain cancers (Gillis et al., 2021; Peixoto et al., 2020). In this regard it must be noted that not all clinically used β-blockers are indicated in all these conditions. For example, only the more lipophilic non-selective molecules which cross the blood brain barrier are used in the treatment of migraine.

Thus far pharmacologists have been limited to understanding β-adrenergic receptor selectivity largely in terms of binding receptor affinity in recombinant systems and in silico structural docking studies that reveal the most favorable, lowest energy binding pose of a ligand (Baker, 2005, 2010; Masureel et al., 2018; Warne et al., 2019; Warne et al., 2011; Warne et al., 2008). However, due to their high sequence and structural similarity the molecular basis of ligand selectivity between these two receptor subtypes remains to be fully understood. This has led to the suggestion that the kinetic process of ligand binding itself may contribute to ligand specificity for individual receptor subtypes. And although the process of ligand binding is not possible to study using direct structural methods, molecular dynamics (MD) studies have shed some light on the molecular basis of receptor selectivity and even on the pathway of ligand binding and dissociation (Dror et al., 2011; Gonzalez et al., 2011; Selvam et al., 2012).

Our previous studies of β_2_-adrenergic receptor kinetics highlighted the importance of ‘membrane like’ polar and hydrophobic interactions in driving ligand binding through changes in ligand association rate (or *k*_on_) (Sykes and Charlton, 2012; Sykes et al., 2014). By itself increased drug affinity for the membrane is unlikely to dictate receptor specific subtype specificity, unless of course a direct lipid entry pathway is involved as has been previously suggested before for the β_2_ -adrenoceptor (Mason et al., 1991). However MD simulation studies appear to rule out a direct lipid pathway over an aqueous route (Dror et al., 2011). Nonetheless, this still does not answer why certain hydrophobic ligands such as salmeterol show preferential binding to the β_2_-adrenoceptor subtype. This appears based on mutation studies to be due partly to the presence of specific residues found on the surface of the β_2_-adrenoceptor which are not found in the corresponding β_1_-adrenoceptor (Baker et al., 2015; Isogaya et al., 1998).

Kinetic selectivity has been shown to be an important factor in dictating the therapeutic action of muscarinic M_3_ antagonists which target the lung to treat chronic obstructive pulmonary disease or COPD (Sykes et al., 2012; Tautermann et al., 2013). Here a longer residency time at the muscarinic M_3_ receptor over the M_2_ receptor subtype is favored from a therapeutic perspective helping to relax smooth muscle cells and open the airways. Blockade of M_2_ receptors results in increased acetylcholine release from vagal nerve endings which enhances bronchoconstriction and precipitates cardiac adverse effects, in particular increases in heart rate, and the induction of polymorphic ventricular tachycardia with the potential for sudden death (Costello et al., 1998; LaCroix et al., 2008; Oberhauser et al., 2001). In direct contrast the relevance of kinetic selectivity in the therapeutic action of β-blockers remains unexplored, largely due to a lack of information on the binding kinetics of these ligands at the β_1_-adrenoceptor.

Historically the kinetics of β_1_-adrenoceptors antagonists have been studied directly in native tissues using tritiated compounds. One of the very earliest examples was formulated in canine myocardium at physiological temperature using tritiated alprenolol, however such experiments are potentially complicated by the presence of multiple receptor subtypes (Alexander et al., 1975). More recently the kinetics of synthetic and endogenous agonists and antagonists have been studied albeit at room temperature (Boursier et al., 2020; Ramos et al., 2018; Xu et al., 2020).

The aim of this work was to determine the kinetics of a series of well-described β_1_-adrenoceptor antagonists and agonists and several clinically relevant β_1_-adrenoceptor antagonists under physiological conditions. One way to achieve this is to indirectly measure the kinetics of drug-receptor interaction using the competition association method first popularized by Motulsky and Mahan (1984) and one we have previously applied with some success to the study of β_2_-adrenergic pharmacology (Sykes and Charlton, 2012; Sykes et al., 2014). A secondary aim was to determine if kinetic selectivity has any role to play in the known side effect profile of these widely prescribed compounds. A more detailed understanding of the molecular basis of β_1/2_-adrenergic receptor antagonist selectivity is likely to provide a novel rationale for the discovery of more selective ligands which target the heart with potentially fewer side effects.

## Materials and Methods

[^3^H]-DHA (1-[4,6-propyl-^3^H]dihydroalprenolol, specific activity 91 Ci·mmol^-1^) was obtained from PerkinElmer Life and Analytical Sciences (Beaconsfield, UK). Ninety-six-deep well plates and 500 cm^2^ cell culture plates were purchased from Fisher Scientific (Loughborough, UK). Millipore 96-well GF/B filter plates were purchased from Receptor Technologies (Warwick, UK). Sodium bicarbonate, ascorbic acid, EDTA, sodium chloride, GTP, bisoprolol hemifumarate, (S)-(-) atenolol, labetalol hydrochloride, (±) metoprolol, carvedilol, (S)-(-) propranolol hydrochloride, salmeterol xinofoate, ICI118,551 hydrochloride, (±) sotalol hydrochloride, nadolol, (S)-(-) cyanopindolol hemifumarate, formoterol fumarate and CGP-20712A methanesulfonate were obtained from Sigma Chemical Co Ltd. (Poole, UK). Bucindolol, (S)-timolol maleate and CGP12177 hydrochloride were obtained from Tocris Cookson, Inc. (Bristol, UK). All cell culture reagents including Hanks’ balanced salt solution (HBSS) and HEPES were purchased from Gibco (Invitrogen, Paisley, UK).

### Cell culture and membrane preparation

CHO cells stably transfected with the human β_1_-adrenoceptor were grown adherently in Ham’s F-12 Nutrient Mix GlutaMAX-1, containing 10% fetal calf serum, and 0.5 mg·mL^-1^ Geneticin (G-418). Cells were maintained at 37°C in 5% CO_2_/humidified air and routinely subcultured at a ratio between 1:10 and 1:20 twice weekly using trypsin-EDTA to lift cells. Cell membranes were prepared and stored as described previously (Sykes & Charlton 2012).

### Common procedures applicable to all radioligand binding experiments

All radioligand binding experiments using 1-[4,6-propyl-^3^H]dihydroalprenolol ([^3^H]-DHA specific activity 91 Ci·mmol-1) were conducted in 96-deep well plates, in assay binding buffer, HBSS pH 7.4, 0.01% ascorbic acid and 100 μM GTP. GTP was included to remove the G protein-coupled population of receptors which can result in two binding sites in membrane preparations, because the Motulsky & Mahan model is only appropriate for ligands competing at a single site. In all cases, nonspecific binding (NSB) was determined in the presence of 1 μM propranolol. After the indicated incubation period, bound and free radiolabels were separated by rapid vacuum filtration using a FilterMate™ Cell Harvester (PerkinElmer Life and Analytical Sciences, Beaconsfield, UK) onto 96 well GF/B filter plates (Millipore, Watford UK) previously coated with 0.5% (w/v) polyethylenimine and rapidly washed three times with ice-cold 75 mM HEPES, pH 7.4. After drying (>4 h), 40 mL of Microscint™ 20 (PerkinElmer Life and Analytical Sciences) was added to each well and radioactivity was quantified using single photon counting on a TopCount™ microplate scintillation counter (PerkinElmer Life and Analytical Sciences). Aliquots of radiolabel were also quantified accurately to determine how much radioactivity was added to each well using liquid scintillation spectrometry on LS 6500 scintillation counter (Beckman Coulter, High Wycombe, UK). In all experiments, total binding never exceeded more than 10% of that added, limiting complications associated with depletion of the free radioligand concentration (Carter et al., 2007).

### Saturation binding studies

CHO cell membranes containing the β_1_-adrenoceptor were incubated in 96-deep well plates at 37°C in assay binding buffer with a range of concentrations of [^3^H]-DHA (∼12 – 0.01 nM) at 30 μg per well, for 180 min with gentle agitation to ensure equilibrium was reached. Saturation binding was performed in a final assay volume of to 1.5 mL to avoid significant ligand depletion.

### Determination of the association rate (k_on_) and dissociation rate (k_off_) of [^3^H]-DHA

To accurately determine *k*_on_ and *k*_off_ values, the observed rate of association (*k*_*ob*_) was calculated using at least three different concentrations of [^3^H]-DHA. The appropriate concentration of radioligand was incubated with β_1_-adrenoceptor CHO cell membranes (30 μg· per well) in assay binding buffer with gentle agitation (final assay volume 500 μL). Exact concentrations were calculated in each experiment by liquid scintillation counting. Free radioligand was separated by rapid filtration at multiple time points to construct association kinetic curves as described previously by Sykes & Charlton (2012). The resulting data were globally fitted to the association kinetic model (Equation 2) to derive a single best fit estimate for *k*_on_ and *k*_off_ as described under Data analysis.

### Determination of affinity constants (K_i_)

To obtain affinity estimates of unlabelled ligand, [^3^H]-DHA competition experiments were performed at equilibrium. [^3^H]-DHA was used at a concentration of approximately 3 nM (final assay volume of 0.5 mL), such that the total binding never exceeded more than 10% of that added. Radioligand was incubated in the presence of the indicated concentration of unlabelled ligand and CHO cell membranes (30 μg per well) at 37°C, with gentle agitation for 180 min.

### Competition binding kinetics

The kinetic parameters of unlabelled ligand were assessed using a competition kinetic binding assay originally described by Motulsky & Mahan (1984) and developed for the β_1_-adrenoceptor by Sykes & Charlton (2012). This approach involves the simultaneous addition of both radioligand and competitor to receptor preparation, so that at t = 0 all receptors are unoccupied.

Approximately 3 nM [^3^H]-DHA (a concentration which avoids ligand depletion in this assay volume) was added simultaneously with the unlabelled compound (at t = 0) to CHO cell membranes containing the human β_1_-adrenoceptor (30 μg· per well) in 500 μL assay buffer. The degree of [^3^H]-DHA bound to the receptor was assessed at several time points by filtration harvesting and liquid scintillation counting, as described previously. NSB was determined as the amount of radioactivity bound to the filters and membrane in the presence of propranolol (1 μM) and was subtracted from each time point, meaning that t = 0 was always equal to zero. Each time point was conducted on the same 96-deep well plate incubated at 37°C with constant agitation. Reactions were considered stopped once the membranes reached the filter, and the first wash was applied within 1 s. A single concentration of unlabeled competitor was tested, as rate parameters were shown to be independent of unlabeled ligand concentration (data not shown). All compounds were tested at either 1-, 3-, 10- or 100-fold their respective *K*_i_ and data were globally fitted using Equation 3 to simultaneously calculate *k*_on_ and *k*_off_. Different ligand concentrations were chosen as compounds with a long residence time equilibrate more slowly so a higher relative concentration is required to ensure the experiments reach equilibrium within a reasonable time frame (90 min), whilst still maintaining a good signal to noise. The actual concentrations used were selected from a preliminary experiment using three different concentrations of each ligand (data not shown).

### LogD_7.4_ and Immobilised Artificial Membrane (IAM) Chromatography

All HPLC experiments were carried out as previously described by Sykes et al., (2014).

#### Data analysis and statistical procedures

As the amount of radioactivity varied slightly for each experiment (<5%), data are shown graphically as the mean ± range for individual representative experiments, whereas all values reported in the text and tables are mean ± SEM for the indicated number of experiments unless otherwise stated. All experiments were analyzed by either Deming regression or non-linear regression using Prism 8.0 (GraphPad Software, San Diego, CA, USA).

### Competition binding

Competition displacement binding data were fitted to sigmoidal (variable slope) curves using a four-parameter logistic equation:

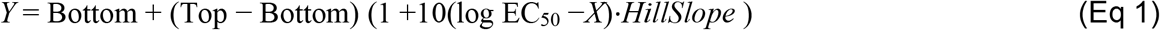

IC_50_ values obtained from the inhibition curves were converted to *K*_i_ values using the method of Cheng and Prusoff (1973).

### Association binding

[^3^H]-DHA association data were globally fitted to Equation 2, where L is the concentration of radioligand in nM using GraphPad Prism 8.0 to determine a best fit estimate for *k*_on_ and *k*_off_.

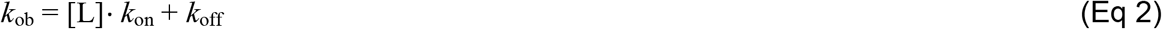

### Competition kinetic binding

Association and dissociation rates for unlabelled agonists were calculated using the equations described by Motulsky and Mahan (1984) using a global fitting model:

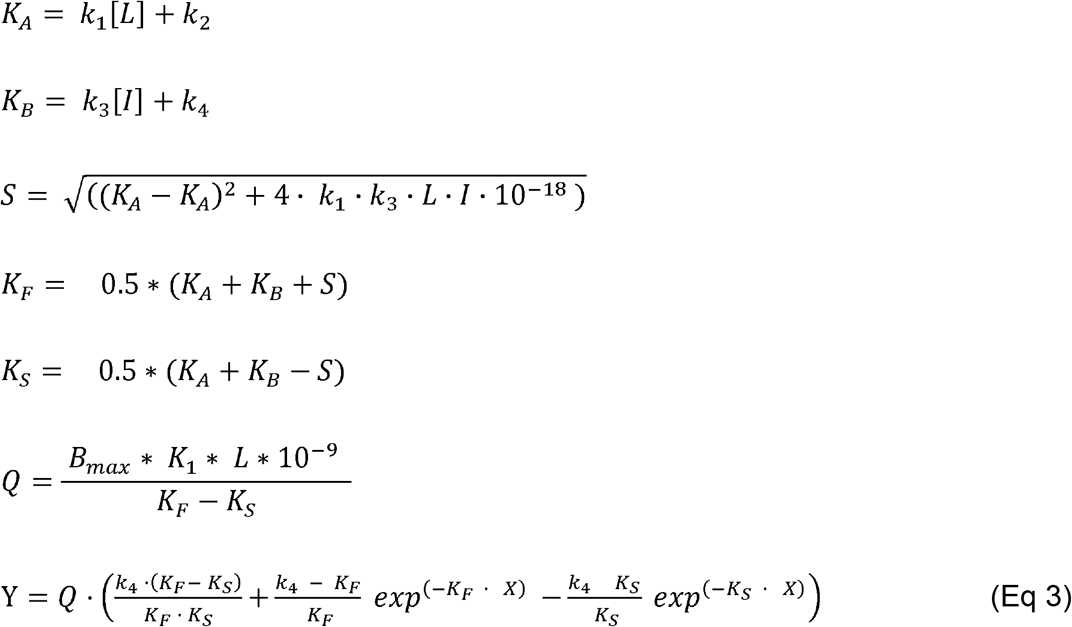

where X is time (min), Y is specific binding (c.p.m.), *k*1 is *k*_on_ of the tracer [^3^H]-DHA, *k*2 is *k*_off_ of the tracer [^3^H]-DHA, *L* is the concentration of [^3^H]-DHA used (nM) and *I* is the concentration of unlabeled agonist (nM). Fixing the above parameters allowed the following to be simultaneously calculated: Bmax is total binding (c.p.m.), *k*3 is association rate of unlabeled ligand (M^-1^ min^-1^) or *k*_on_, and *k*4 is the dissociation rate of unlabelled ligand (min^-1^) or *k*_off_.

### Linear correlations

The correlation between datasets was determined by calculating a Pearson correlation coefficient (presented as r^2^ the coefficient of determination, which shows percentage variation in y which is explained by all the x variables together) in GraphPad Prism 8.0.

#### PDBs structures and hydrophobic and electrostatic surface maps

The crystal structures of the β_1_AR (Xu et al., 2020) and β_2_AR (Staus et al., 2016) were obtained from the Protein Data Bank (PDB entries 7BVQ and 5JQH, respectively). The hydrophobic surface map was obtained using a modified version (changing the scale of colours to purple and yellow) of color_h.py script (pymolwiki.org/index.php/Color_h) using PyMOL. For production of the electrostatic surface map, we used the PyMOL APBS electrostatic plugin using the default parameters (changing the scale colours to blue and red only) (MG Lerner and HA Carlos. APBS plugin for PyMOL, 2006, University of Michigan, Ann Arbor).

## Results

### Equilibrium and kinetic binding parameters for β_1_-adrenoceptor ligands

Initially the binding affinity of the radioligand [^3^H]-DHA for the β_1_-adrenoceptor (shown in Figure 1A) was measured at equilibrium in Hanks’ Balanced Salt Solution containing GTP (100μM) at 37°C. GTP was included to ensure that agonist binding only occurred to the uncoupled form of the receptor. Binding affinities (*K*_i_ values) for the β_1_-adrenoceptor ligands determined in the presence to GTP are summarized in Table 1 and associated curves are presented in Figure 1B-D. To determine the association and dissociation rates of the β_1_-adrenoceptor ligands, we utilized a competition kinetic radioligand binding assay as previously described by Sykes & Charlton (2012). Firstly, we characterized the binding kinetics of the radiolabelled ligand [^3^H]-DHA by monitoring the observed association rates at 3 different ligand concentrations (Figure 2A). The observed rate of association was related to [^3^H]-DHA concentration in a linear fashion (Figure 2B). Kinetic rate parameters for [^3^H]-DHA were calculated by globally fitting the association time courses, resulting in a *k*_on_ of 5.28 ± 0.48 ×10^9^ M^-1^ min^-1^ and a *k*_off_ of 0.46 ± 0.03 min^-1^ (*K*_d_ = 0.94 ± 17 nM).

**Table 1.** Kinetic binding parameters of unlabeled ligands for human β_1_-adrenoceptor receptors. Data are mean ± SEM for ≥3 experiments performed in singlet.

| Ligand | $k_{\text{off}}$ ($\text{min}^{-1}$) | $k_{\text{on}}$ ($\text{M}^{-1} \text{min}^{-1}$) | Residence time (min) | $\text{pK}_d$ | $\text{pK}_i$ |
| --- | --- | --- | --- | --- | --- |
| Bisoprolol | $0.14 \pm 0.06$ | $1.96 \pm 0.26 \times 10^7$ | 7.14 | $8.20 \pm 0.13$ | $7.86 \pm 0.11$ |
| Atenolol | $6.19 \pm 2.38$ | $4.03 \pm 1.29 \times 10^7$ | 0.16 | $6.84 \pm 0.04$ | $6.59 \pm 0.22$ |
| Labetolol | $1.74 \pm 0.55$ | $2.85 \pm 0.91 \times 10^8$ | 0.57 | $8.21 \pm 0.07$ | $8.01 \pm 0.12$ |
| Metoprolol | $3.41 \pm 0.85$ | $1.24 \pm 0.11 \times 10^8$ | 0.29 | $7.59 \pm 0.10$ | $7.20 \pm 0.15$ |
| Bucindolol | $0.11 \pm 0.02$ | $1.11 \pm 0.25 \times 10^9$ | 9.09 | $10.00 \pm 0.04$ | $9.57 \pm 0.11$ |
| Carvedilol | $0.09 \pm 0.02$ | $1.97 \pm 0.39 \times 10^9$ | 11.11 | $10.36 \pm 0.13$ | $9.84 \pm 0.23$ |
| Propranolol | $0.91 \pm 0.03$ | $1.28 \pm 0.22 \times 10^9$ | 1.10 | $9.13 \pm 0.09$ | $9.51 \pm 0.09$ |
| Salmeterol | $14.3 \pm 3.65$ | $2.63 \pm 0.21 \times 10^7$ | 0.07 | $6.29 \pm 0.08$ | $5.92 \pm 0.05$ |
| ICI 118, 551 | $7.40 \pm 1.31$ | $6.80 \pm 2.40 \times 10^7$ | 0.14 | $6.88 \pm 0.16$ | $6.14 \pm 0.13$ |
| Sotalol | $8.49 \pm 3.23$ | $8.47 \pm 3.16 \times 10^6$ | 0.12 | $6.00 \pm 0.03$ | $5.60 \pm 0.05$ |
| Nadolol | $0.17 \pm 0.03$ | $7.68 \pm 0.79 \times 10^6$ | 5.88 | $7.65 \pm 0.04$ | $7.47 \pm 0.15$ |
| Formoterol | $9.97 \pm 2.43$ | $1.68 \pm 0.11 \times 10^7$ | 0.10 | $6.25 \pm 0.12$ | $5.84 \pm 0.05$ |
| Timolol | $0.05 \pm 0.01$ | $9.66 \pm 2.42 \times 10^7$ | 20.00 | $9.24 \pm 0.06$ | $8.75 \pm 0.05$ |
| Cyanopindolol | $0.015 \pm 0.003$ | $2.99 \pm 0.23 \times 10^8$ | 66.67 | $10.34 \pm 0.09$ | $9.93 \pm 0.09$ |
| CGP12177 | $0.08 \pm 0.02$ | $5.07 \pm 0.93 \times 10^8$ | 12.50 | $9.83 \pm 0.11$ | $9.50 \pm 0.22$ |
| Isoprenaline | $4.77 \pm 1.74$ | $2.51 \pm 0.44 \times 10^7$ | 0.21 | $6.78 \pm 0.16$ | $6.49 \pm 0.09$ |
| Salbutamol | $9.81 \pm 2.60$ | $1.96 \pm 0.77 \times 10^7$ | 0.10 | $4.91 \pm 0.02$ | $5.27 \pm 0.11$ |
| CGP20712A | $0.22 \pm 0.03$ | $4.14 \pm 0.60 \times 10^8$ | 4.55 | $9.27 \pm 0.08$ | $8.75 \pm 0.06$ |

**Figure 1.**
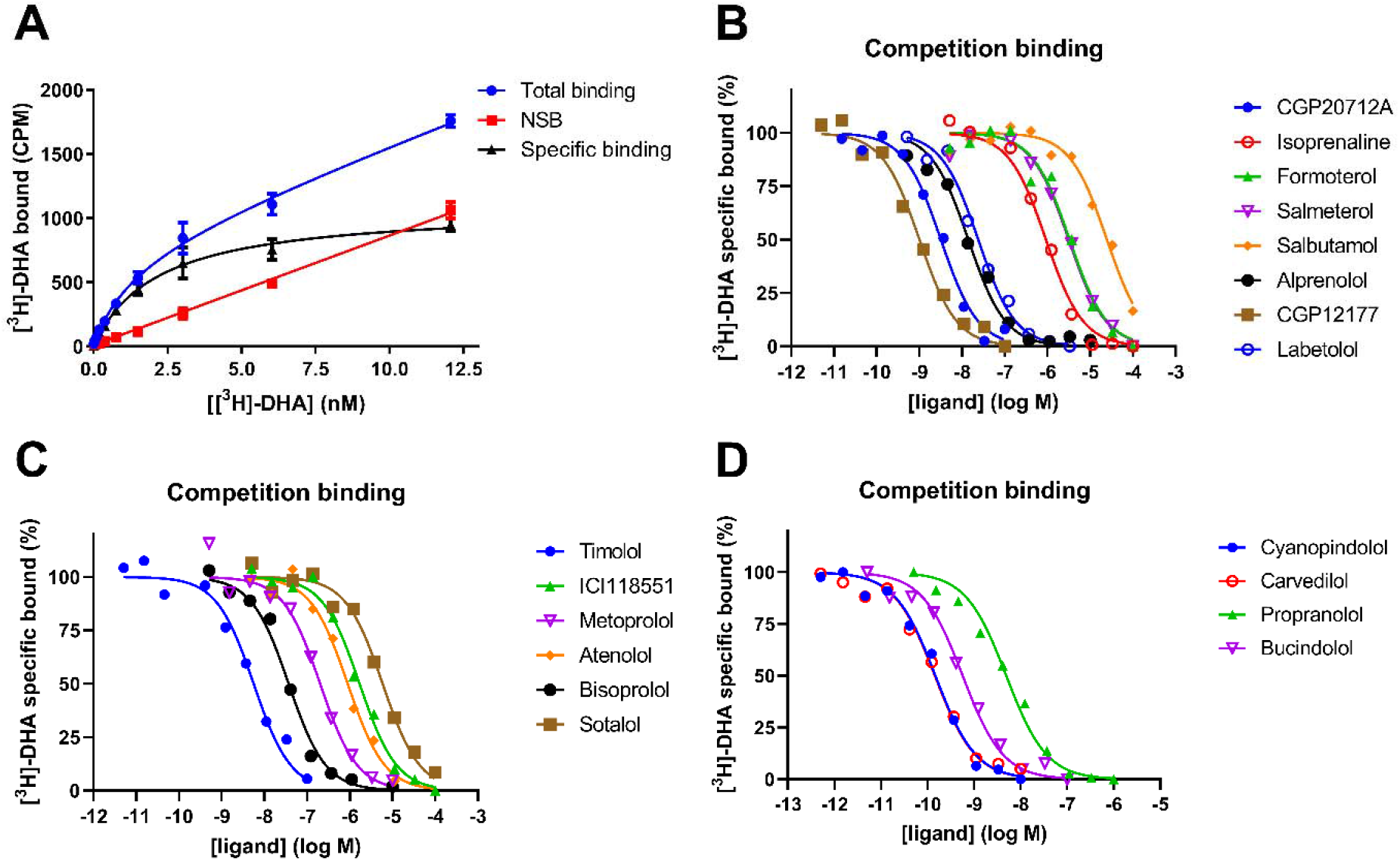
[^3^H]-DHA saturation binding plus competition binding between [^3^H]-DHA and β-adrenergic ligands for human β_1_-adrenoceptors expressed in the CHO cells in the presence of GTP. **(A)** [^3^H]-DHA saturation binding, Membranes (30 μg per well) from CHO-β1 cells were incubated in HBSS containing 0.1 mM GTP (as described in Methods) and increasing concentrations of radioligand for 180min at 37°C with gentle agitation. Displacement of [^3^H]-DHA (3nM) by increasing concentrations of **(B)** CGP20712A, isoprenaline, formoterol, salmeterol, salbutamol, alprenolol, CGP12177 and labetolol. **(C)** timolol, ICI118,551, metoprolol, atenolol, bisoprolol, sotalol. **(D)** (S)-cyanopindolol, carvedilol, propranolol and bucindolol. NSB was defined by 1 μM propranolol. Data are presented as the mean ± range from a representative of 3 experiments performed in singlet.

**Figure 2.**
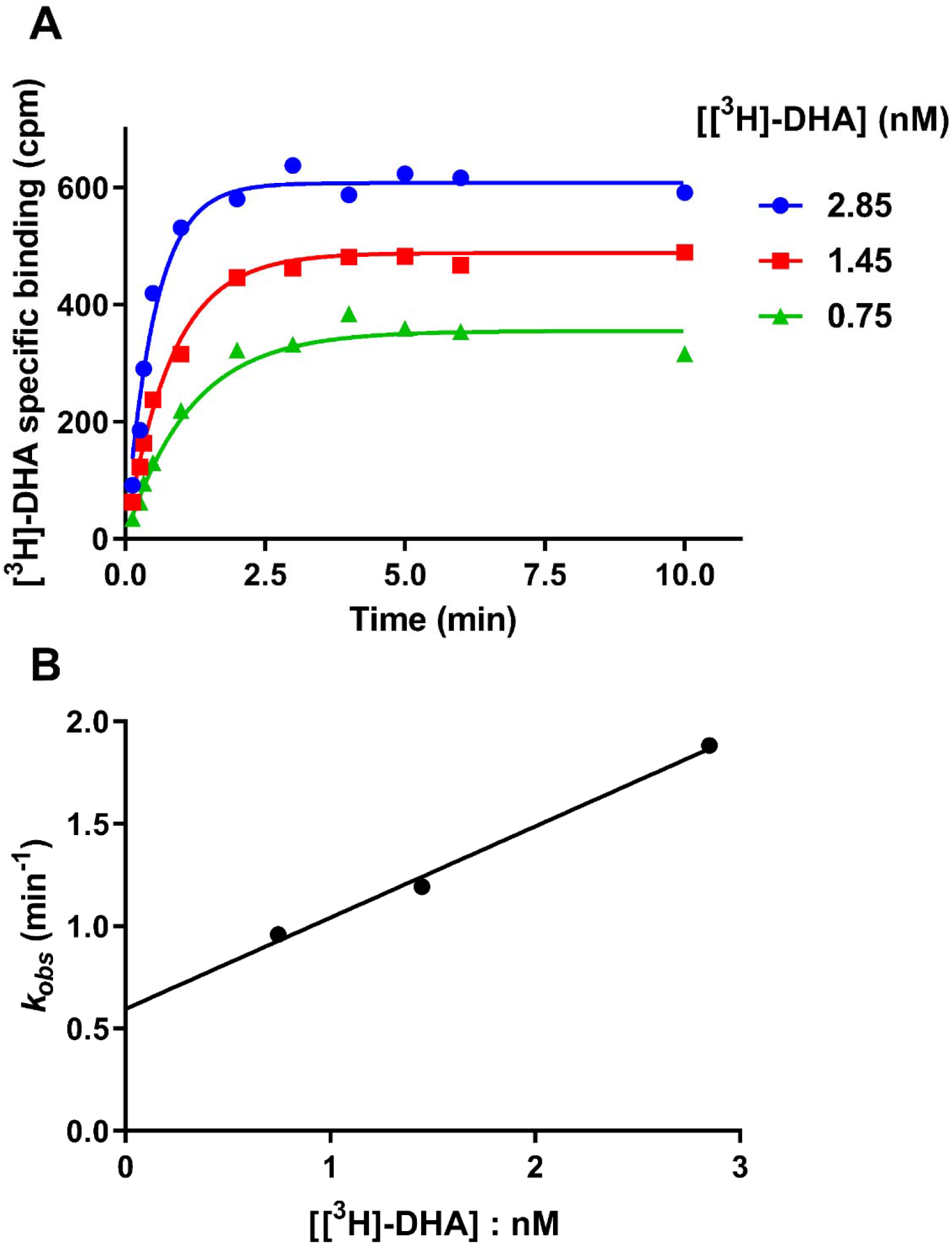
Determination of [^3^H]-DHA kinetic binding parameters. **(A)** Observed association kinetics of the interaction of [^3^H]-DHA with CHO membranes expressing the human β_1_-adrenoceptors. **(B)** Plot of ligand concentration verses *k*_ob_. Binding followed a simple law of mass action model, *k*_ob_ increasing in a linear manner with radioligand concentration. Data are presented as the mean ± range from a representative of 3 experiments performed in singlet.

Representative kinetic competition curves for selected β-adrenoceptor ligands are shown in Figure 3A-H. Progression curves for [^3^H]-DHA alone and in the presence of competitor were globally fitted to Equation 3 enabling the calculation of both *k*_on_ (*k*3) and *k*_off_ (*k*4) for each of the ligands, as reported in Table 1. There was a very wide range in dissociation rates for the different ligands, with drug-target residency times (1/*k*_off_) ranging between 0.07 min for the rapidly dissociating salmeterol to 66.67 min for cyanopindolol. To validate the rate constants, the kinetically derived *K*_d_ values (*k*_off_/*k*_on_) were compared with the affinity constant (*K*_i_) obtained from equilibrium competition binding experiments (Supplemental Figure 1). There was a very good correlation (*r*^2^ = 0.97, *P* < 0.0001) between these two values, indicating the kinetics parameters were accurate.

**Figure 3.**
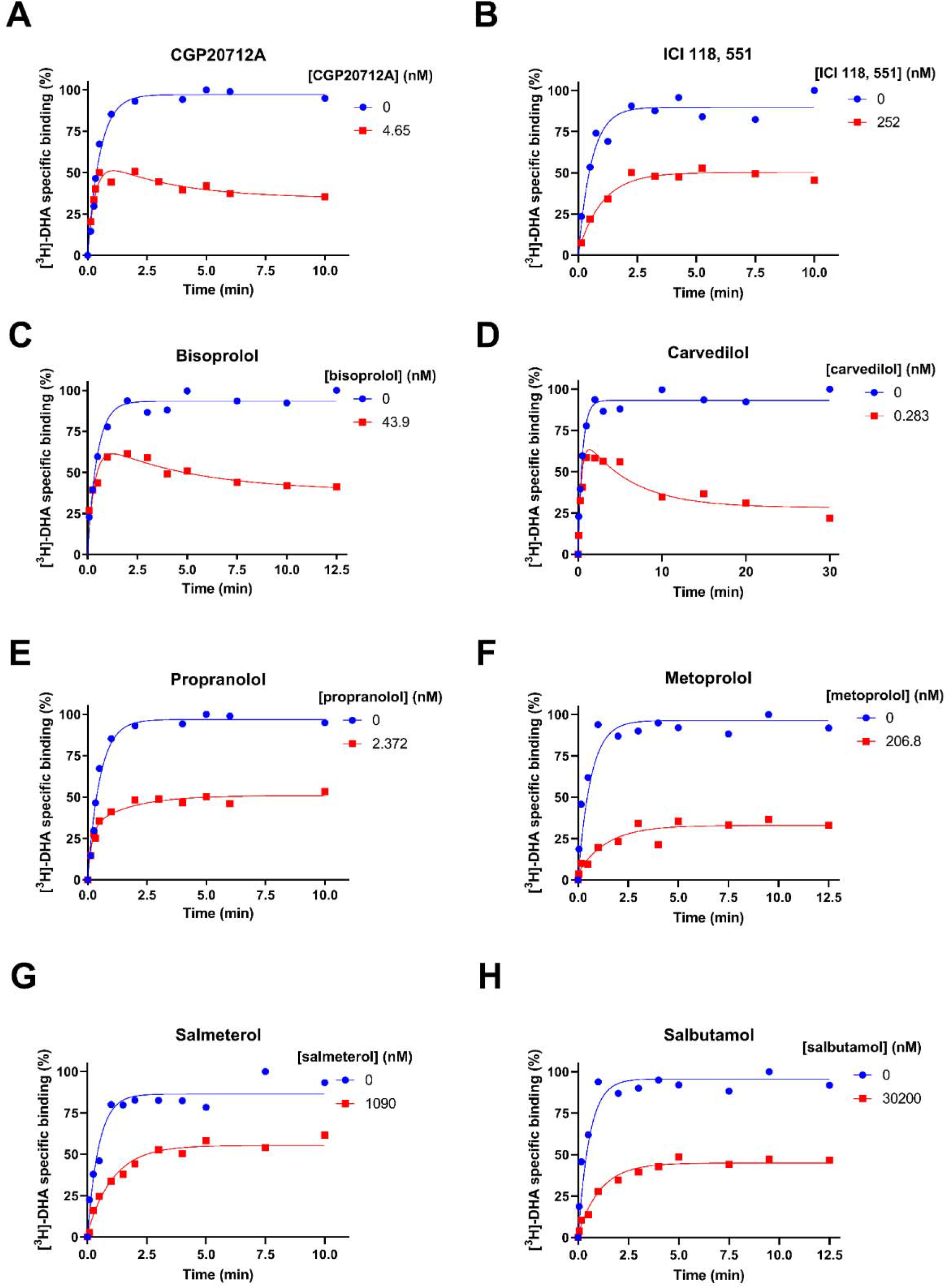
[^3^H]-DHA competition kinetic curves in the presence of CGP20712A **(A)**, ICI118,551**(B)**, bisoprolol **(C)**, carvedilol **(D)**. propranolol **(E)** metoprolol **(F)** salmeterol and **(H)** salbutamol. CHO-β_1_ membranes were incubated with ∼3 nM [^3^H]-DHA and either 0-, 1-, 3-, 10- or 100-fold *K*_i_ of unlabeled competitor. Plates were incubated at 37°C for the indicated time points and NSB levels were determined in the presence of 1 μM propranolol. Data were fitted to the equations described in the Methods to calculate *k*_on_ and *k*_off_ values for the unlabelled ligands; these are summarized in Table 1. Data are presented as mean ± range from a representative of ≥3 experiments performed in singlet.

### Measurements of lipophilicity and membrane interactions

The degree of membrane interaction, denoted *K*_IAM_, assessed using a chromatographic method and calculated logP (clogP) values and the measured partition coefficient logD_7.4_ are detailed in (Sykes et al., 2014). Drug membrane interaction is assessed using a chromatographic method that utilises immobilized artificial membranes (IAMs) consisting of monolayers of phospholipid covalently immobilized on a silica surface, mimicking the lipid environment of a fluid cell membrane on a solid matrix (Liu et al., 2001; Sykes et al., 2014). Compounds with longer retention times on this column have higher affinity for phospholipids and will therefore theoretically have a higher calculated membrane partition coefficient, denoted *K*_IAM_. The main difference between the logD_7.4_ measure, that reflects hydrophobicity of the compound and IAM systems being the key role of electrostatics in the differential binding of the charged ligands to the anisotropic IAM column (Vrakas et al., 2008), mimicking the effects seen in biological membranes.

### Relationship between kinetics and membrane interactions

The association rate parameter *k*_on_ is calculated from the observed on-rate (*k*_obs_) which is highly dependent on drug concentration. We have demonstrated previously that drugs possessing a high membrane affinity appear to have a more rapid association rate, potentially due to an increase in the local concentration of compound (Sykes et al., 2014; Gherbi et al., 2017).

The degree of membrane interaction is routinely assessed using a chromatographic method. The calculated membrane partition coefficient, denoted *K*_IAM_, logP (clogP) and the measured partition coefficient logD_7.4_ used in the following plots are detailed in Sykes et al., 2014. Test compound Log D_7.4_ values were significantly correlated (*r*^*2*^ = 0.32, *P*=0.019) with *k*_on_ determined in the competition kinetic assay (Figure 4A), with lipophilic compounds having a faster association rate. A very similar correlation of *k*_on_ with log*K*_IAM_ was observed (Figure 4B, *r*^*2*^ = 0.35, *P*=0.013), suggesting that a simple isotropic, single parameter model may be sufficient to describe the interaction of these drugs with the β_1_-adrenoceptor. This was supported by comparisons between clogP and *k*_on_ which also showed a better correlation than observed with K_IAM_ (*r*^*2*^ = 0.48, *P*<0.002, data not shown). CGP20712A was excluded from these and subsequent analysis as no *K*_IAM_ data or corresponding β_2_AR binding data was available.

**Figure 4.**
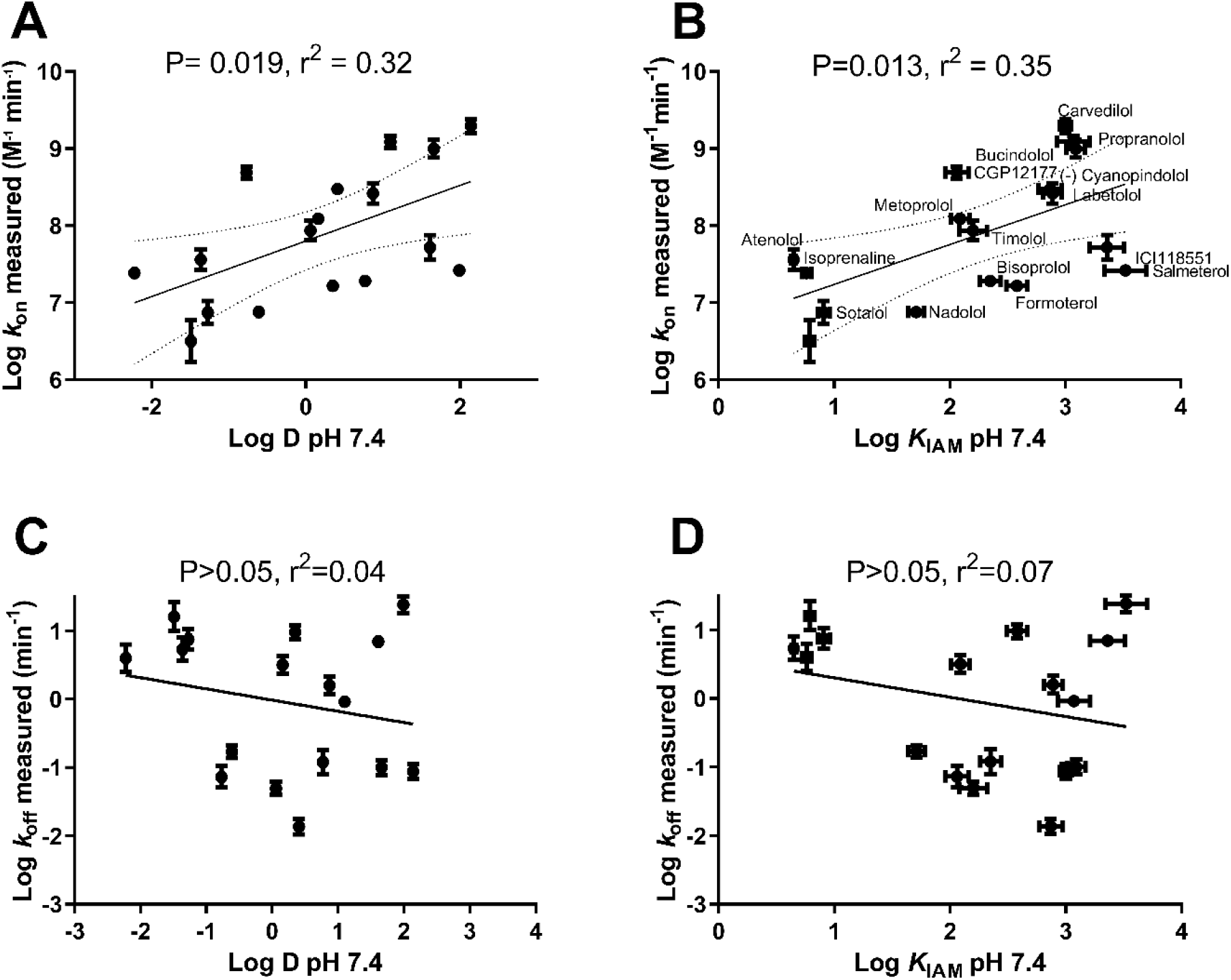
Correlating β_1_ adrenoceptor ligand physiochemical parameters with kinetically derived parameters. Correlation plot showing the relationship between **(A)** log *k*_on_ and log D_7.4_ and **(B)** log *k*_on_ and log K_IAM7.4._ Correlation plot showing the relationship between **(C)** log *k*_off_ and log D_7.4_ and **(D)** log *k*_off_ and log K_IAM7.4_. All data used in these plots are detailed in Table 1 with log D_7.4_ and log K_IAM7.4_ values from Sykes et al., 2014. Data are presented as mean ± SEM from three or more experiments.

The dissociation rate (or *k*_off_) of a drug is not dependent upon drug concentration, so should be independent of the affinity of interaction with the membrane. Reassuringly, when the *k*_off_ for each compound was compared to either its logD_7.4_ or log*K*_IAM_ no correlation was observed (P>0.05, Figures 4C and 4D respectively).

The role of kinetics in dictating β_1_-adrenoceptor compound affinity is presented in Figure 5A. Of the clinically used β-blockers under study, bisoprolol and nadolol stand out as possessing relatively slow on-rates in the region of ∼10^7^ M^-1^min^-1^ and relatively slow off rates (∼0.1min^-1^). In contrast other clinically used agents such as metoprolol and atenolol have much faster dissociation rates (3-10min^-1^) but higher relative association rates.

**Figure 5.**
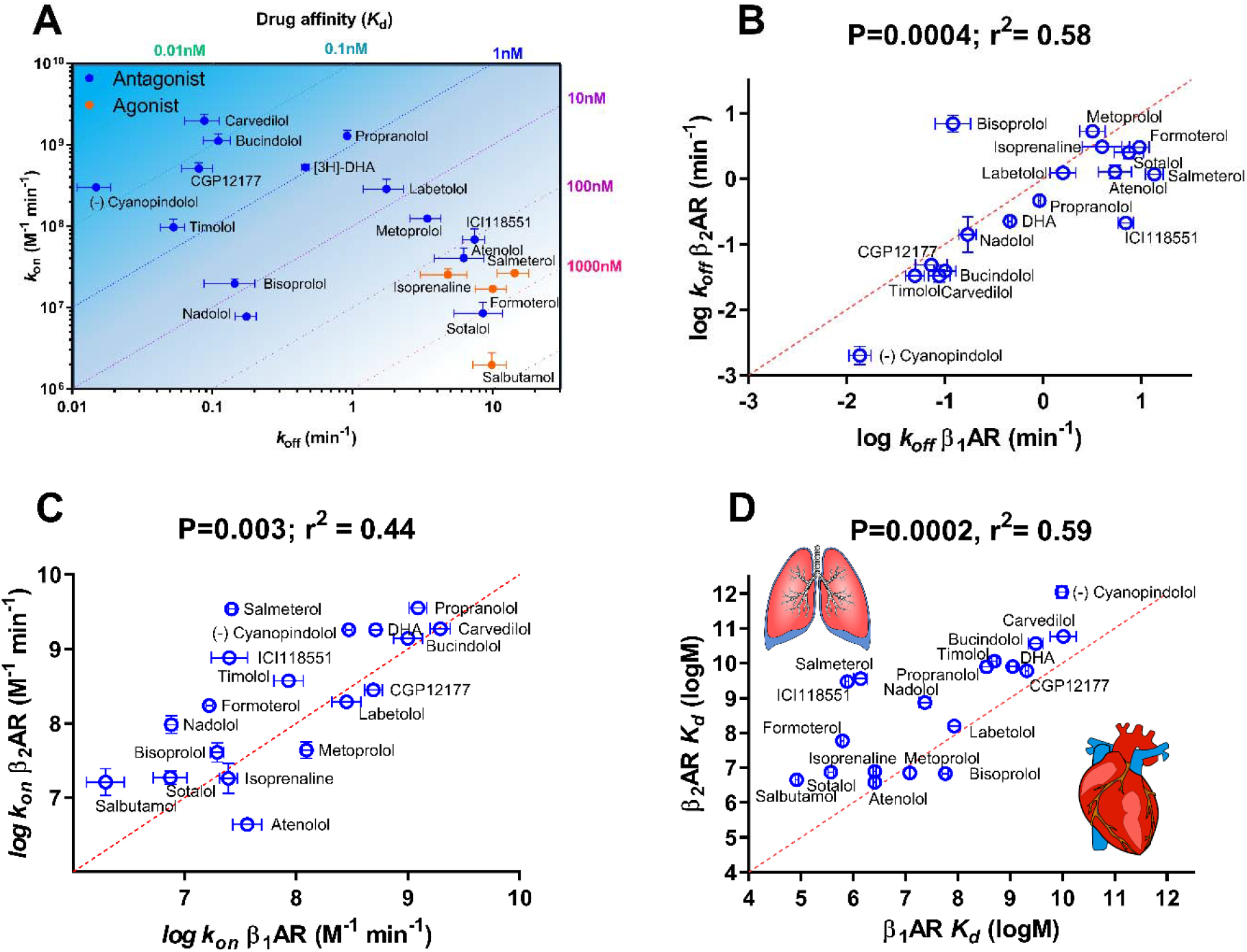
Summarizing the role of kinetics in dictating β_1_-adrenoceptor affinity and selectivity. **(A)** Plot of β_1_AR *k*_off_ versus β_1_AR *k*_on_ values with affinity indicated by the diagonal dotted lines. **(B)** Plot of β_1_AR *k*_off_ versus β_2_AR *k*_off_ values. **(C)** Plot of β_1_AR *k*_on_ versus β_2_AR *k*_on_ values. **(D)** Plot of β_1_AR log *K*_d_ versus β_2_AR log *K*_d_ values. Kinetic values are presented as mean ± SEM from three or more experiments detailed in Table 1.

A comparison β_1/2_-adrenoceptor compound dissociation and association kinetics is presented in Figure 5B to C. Of the clinically used β-blockers bisoprolol again stands out as the only truly β_1_-adrenoceptor selective compound based on its kinetic affinity, a feature that is seemingly dictated by its dissociation rate from the β_1_-adrenoceptor (see Figure 5B and D). The majority of ligands demonstrate a faster association rate at the β_2_-adrenoceptor apart from the clinically used β-blocker atenolol (Figure 5C). Other key observations in terms of understanding β-adrenergic ligand selectivity are the pronounced reduction in the β_1_-adrenoceptor association and dissociation rate of salmeterol relative to salbutamol when we compare kinetic values across the two receptors (Figure 5B and C). Similarly, ICI 118, 551 lower affinity for the β_1_-adrenoceptor appears to be dictated by a combination of a reduced association and increased dissociation rate.

### Using kinetic parameters to model the rate of receptor occupancy and dissociation from the β_1_ and β_2_ adrenoceptors

The rate of receptor occupancy is one factor which could potentially play a significant role in the rate of onset of the actions of clinically used β-blockers. To investigate this, we stimulated their *k*_obs_ at the β_1_ and β_2_ adrenoceptors using a concentration 30\**K*_d_ their β_1_-adrenoceptor affinity. Under these conditions there were clear differences in the rate of association of the four clinically used compounds with bisoprolol and carvedilol exhibiting a slower rate of β_1/2_ receptor occupancy than the other clinically used ligands tested metoprolol and atenolol (Figure 6A-D) that saturated the receptors faster.

**Figure 6.**
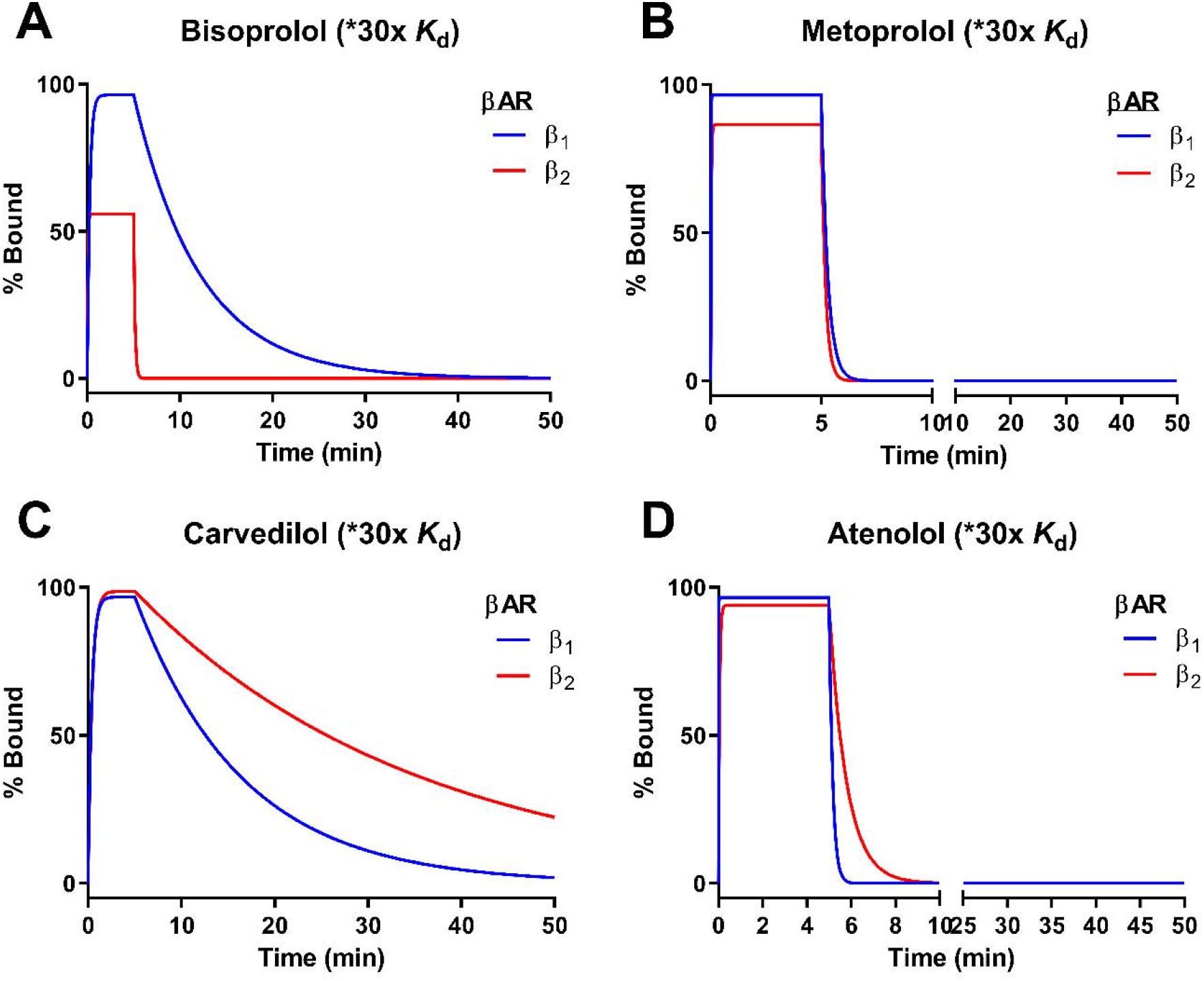
Summarizing the role of kinetics in dictating β_1_-adrenoceptor selectivity. Modelling the association and dissociation of clinically relevant β-blockers. Simulated binding of clinically relevant β-blockers **(A)** bisoprolol **(B)** metoprolol **(C)** carvedilol and **(D)** atenolol to human β_1_-adrenoceptors and β_2_-adrenoceptors at concentrations 30\**K*_d_ of the β_1_-adrenoceptor.. Dissociation occurs at 5 min point initiated by the removal of free ligand. The kinetic parameters used to construct these simulations are detailed in Table 1

Another factor which could play a role in the duration of action (DOA) of clinically used β-blockers is their rates of dissociation from the β-adrenoceptors. In the simulations compound dissociation is initiated by the removal of free ligand at the 5 min mark. These simulations show that for atenolol and metoprolol receptor binding is fully reversed within 5 min, suggesting that dissociation rate has little or no role to play in the DOA of these two clinically used compounds (Figure 6B and D). Atenolol achieves a marginally lower level of occupancy at the β_2_-adrenoceptors but its slower dissociation rate from this receptor equates to a marginally extended occupancy at this receptor.

In contrast dissociation of bisoprolol is noticeably slower from the β_1_ -adrenoceptors compared to the β_2_ -adrenoceptor. And whilst full dissociation occurs from the β_2_ - adrenoceptor in a matter of seconds, it takes approximately 40 min for full dissociation from the β_2_ -adrenoceptor (Figure 6A). Carvedilol is a 3^rd^ generation high affinity β-blocker and it is noticeable that it displays both higher affinity for the β_2_-adrenoceptor but also a much slower rate of dissociation which leads to an extended occupancy at this receptor (Figure 6C). β_1_-adrenoceptors residence time values from these simulations are detailed in Table 1. Residence time values for β_2_-adrenoceptors were taken from a previous publication (Sykes et al., 2014).

## Discussion

This study reports the kinetic rate constants of a number of β_1_-adrenoceptor antagonists and agonists under physiological conditions allowing direct comparisons with earlier kinetic studies of the β_2_-adrenoceptor (Sykes and Charlton, 2012; Sykes et al., 2014). Previous findings demonstrate how local drug concentrations near receptors embedded in biological membranes can directly influence their observed pharmacology (Gherbi et al., 2018; Sykes et al., 2014). Having established that a membrane bilayer acts as a medium by which drug molecules interact or locate low concentrations of a receptor, we proposed that compounds with high membrane partitioning would result in increased values of *k*_on_. We have now extended these observations to include the β_1_-adrenoceptor, comparing observed kinetic rate parameters with the degree of interaction with immobilized artificial membranes (K_IAM_) and measures of lipophilicity (Log P and logD_7.4_). As predicted the association rate of the compounds was seemingly directly influenced by their lipophilicity (logD_7.4_) however surprisingly the magnitude of interaction with the membrane surrounding the receptor, as determined through the artificial membrane partition coefficient (or K_IAM_), did not further enhance this correlation.

In the previous study of the β_2_-adrenoceptor, we hypothesized that the membrane itself could interact specifically with drugs (Sargent and Schwyzer, 1986), through ionic and hydrogen-bonding interactions (Avdeef et al., 1998), effectively concentrating drug molecules close to the surface of the membrane (relative to the bulk solution). In addition to the drug concentrating effect of the membrane, the loss of drug associated water (Dror et al., 2011), and lateral diffusion across a 2-dimensional cellular surface (rather than 3-dimensions in aqueous bulk) could all contribute to increased ligand-receptor association rates (McCloskey and Poo, 1986). These rate enhancing effects may not only be applied to membrane-like structures (e.g., phospholipids) but also the extracellular surfaces of the receptor itself (e.g., amino acids with a polar, hydrophilic and a nonpolar, hydrophobic end). A compound can have the right physicochemical properties to facilitate a fast on-rate but it must also have the right complementary structural features to facilitate its interaction with the receptors binding pocket.

Mutational, crystal modelling and docking studies have highlighted the key role that specific regions of these receptors forming the entrance to the binding pocket, play in dictating overall drug-receptor affinity (Baker et al., 2015; Dror et al., 2011; Isogaya et al., 1999; Kaszuba et al., 2010; Kikkawa et al., 1998; Plazinska et al., 2013; Plazinska et al., 2015; Warne et al., 2008) and kinetics (Xu et al., 2020). A comparison of the residues of both β-adrenoceptor subtypes has suggested the importance of non-conserved electrostatic interactions as well as conserved aromatic contacts in the early steps of the binding process (Selvam et al., 2012; Xu et al., 2020). Similarly, on exit molecules have been shown to pause in what would now be termed the extracellular vestibule, a site 9-15 Å from the orthosteric binding site. These same sites have been shown to serve as secondary binding pockets during MD simulations of ligand entry (Dror et al., 2011; Gonzalez et al., 2011).

Based on the above one plausible explanation for differences in drug-receptor subtype association rate stems from the different amino acid composition of their vestibular regions (Masureel et al., 2018; Vanni et al., 2009; Warne et al., 2008). The extracellular vestibule of the β_2_AR is well known to have a more extensive polar network (Ring et al., 2013; Xu et al., 2020). In the β_2_-adrenoceptor structure, the polar hydrophobic amino acid Tyr308^7.35×34^ contains an oxygen atom in the side chain which is capable of acting as a H-bond donor or acceptor and has been postulated to form a key node in the pathway to successful ligand binding (Dror et al., 2011). Superscript notation corresponds to the GPCRDB numbering (Isberg et al., 2016). Mutation of this residue to structurally equivalent phenylalanine found in the β_1_-adrenoceptor structure (Phe259^7.35×34^), and a purely hydrophobic amino acid, has been shown to reduce the binding affinity of a wide range of β_2_ adrenergic ligands firmly establishing its importance in the binding process (Baker et al., 2015). The absence of this key polar residue in the β_1_-adrenoceptor structure means that for certain ligands the connection between their ability to interact with the membrane and subsequent probability of a successful binding event is reduced. The idea that specific residues (hydrophobic/charged) residues may guide lipophilic polar molecules into the orthosteric binding pocket, it is very much in line with predictions from MD studies which highlight the probability of binding from different regions of the outer pocket (Dror et al., 2011). This concept is also consistent with the idea that the receptor itself can influence local drug concentrations and in so doing directly dictate drug-receptor affinity through an increase in measured on-rate (Gherbi et al., 2018). The relative differences in extracellular surface lipophilicity and polarity between the β_1_ and β_2_-adrenoceptor subtypes are summarized in Figure 7.

**Figure 7.**
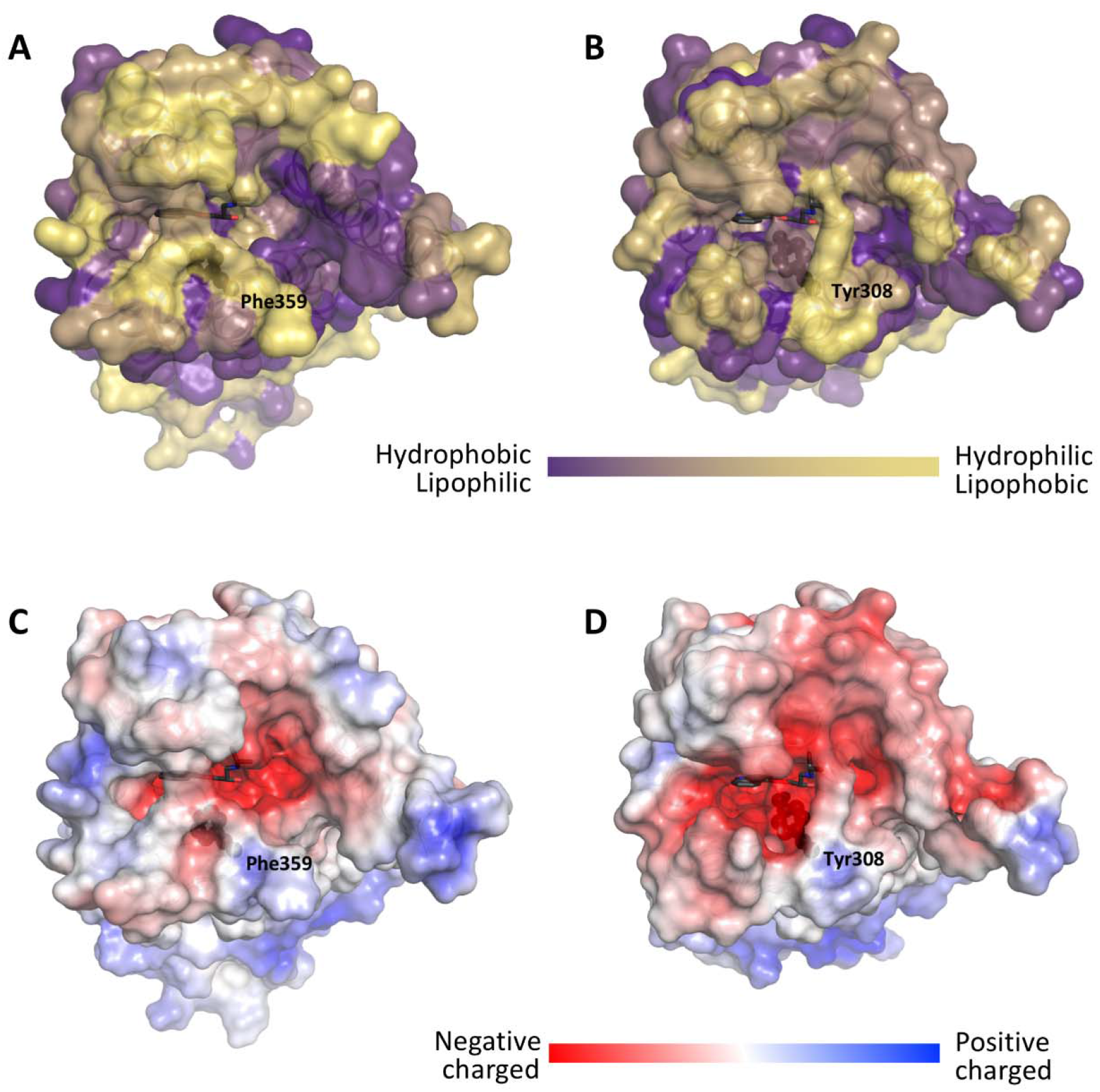
Comparison of the extracellular surfaces including vestibules and orthosteric pockets between the human β_1_AR and β_2_AR. Extracellular surface lipophilicity of the β_1_AR and β_2_AR is shown in **(A)** and **(B)** respectively. Extracellular surface polarity of the β_1_AR and β_2_AR is shown in **(C)** and **(D)** respectively. Overall, the surface of the β_2_AR is dominated by more negatively charged patches and its hydrophobic regions are more numerous and widely spread.

Ligand binding likely occurs in a multistage process: Initially the membrane acts as a vehicle to concentrate drug molecules at the receptor surface contributing to the loss of water molecules and allowing binding to proceed in several ‘smaller’ steps, with lower energy barriers compared with a one-step mechanism, culminating in a more rapid binding process (AP and Guo, 2019; Vauquelin, 2016). Diffusion in 2D allows drugs molecules to rapidly reach their target receptor through a reduction in dimensionality. Drug then enters the vestibule, followed by interaction of a ligand with key residues in the outer vestibule, for example in the β_2_-adrenoceptor structure certain basic drug molecules form a loose interaction with the more numerous polar amino acid residues eg. Tyr308^7.35×34^ (Y), thereby increasing the probability of a successful drug binding event. Finally, as a ligand enters the negatively charged binding pocket the positive charge of the ligands contributes to their rapid association with the orthosteric binding site.

It is reasonable to assume that the process of unbinding and binding follows a similar path but in opposite directions. A slower dissociation rate from the β_2_-adrenoceptor could be partly caused by an extended residence time in the vestibule of receptor, the result of both the polar and hydrophobic interactions important in facilitating ligand association. This being the case the absence of Tyr308^7.35×34^ in the β_1_-adrenoceptor structure should result in an overall more rapid dissociation rate for key β_2_-adrenoceptor specific ligands. This idea appears to be consistent with both the kinetic on and off-rates measured at these two receptor subtypes (Ramos et al., 2018; Sykes et al., 2012; Sykes et al., 2014) and the reduced affinities observed for ligands following the Y308F mutation (Baker et al., 2015). The relationship between association and dissociation rates for these ligands across these two receptor subtypes is shown in Supplementary Figure 2.

Kinetic selectivity is likely to be one of the key steps to reducing the side effect profile of β-adrenergic compounds. This tactic has proved to be effective in reducing the side effect profile of muscarinic M_3_ antagonists which based on affinity values cannot be considered particularly selective for one receptor subtype over another (Sykes et al., 2012; Tautermann et al., 2013). In the current study, we are able for the first time to rationalize the improved affinity of bisoprolol for the β_1_-adrenoceptor over the β_2_-adrenoceptor which results from its much-reduced dissociation rate (50-fold, see Figure 5b and 6a) (Sykes et al., 2014).

Bisoprolol’s use in the clinic in patients with chronic heart failure and COPD is recognized as being associated with fewer side effects and potentially an improvement in mortality compared to the other β-blockers a fact which could be rationalized based on its improved kinetic selectivity for the β_1_-adrenoreceptors, cumulating in fewer off-target effects and characterized by its slow elimination and accumulation in the heart (Kang et al., 2021; Kubota et al., 2015; Lainscak et al., 2011; Liao et al., 2017; Su et al., 2016; Taniguchi et al., 2013). Other clinically used β-blockers which are considered cardio-selective include metoprolol and atenolol. However, based on the current data only atenolol can be considered marginally selective in terms of its overall affinity for β_1_ and β_2_-adrenoceptors. Neither drug can be considered kinetically selective in terms of their measured off-rates. Other non-selective β_2_-adrenoceptor blockers such as carvedilol appear to be if anything more selective for β_2_-adrenoreceptors over β_1_-adrenoreceptors with repercussions for patients with underlying respiratory disease (Baker and Wilcox, 2017; Benson et al., 1978) where greater reductions in FEV (lung function) have been observed (Dungen et al., 2011; Jabbal et al., 2017; Jabbour et al., 2010) (see Figure 6).

Bisoprolol selective effects in the heart are clearly beneficial in the patients with heart and lung disease but in terms of the treatment of heart failure some reports suggest that carvedilol may produce equivalent or an improved overall reduced chance of all-cause mortality in systolic heart failure compared to other more selective agents such as bisoprolol (Bølling et al., 2014; Choi et al., 2019; DiNicolantonio et al., 2013; Hart, 2000; Hulkower et al., 2015; Rain and Rada, 2015; Remme, 2010; Wikstrand et al., 2014). Any number of factors could contribute to this including carvedilol’s relatively slower dissociation from the β_1_-adrenoreceptor or its vasodilatory alpha-receptor blocking effects (Metra et al., 2004; Remme, 2010; Weir and Dargie, 2005). Alternatively, its apparent kinetic selectivity for the β_2_-adrenoreceptor, and/or biased signaling profile could potentially contribute to beneficial remodeling effects in the heart (Kim et al., 2014; Wisler et al., 2007).

In conclusion, we hope that the new kinetic data outlined in this study will reignite research into the discovery and development of kinetically selective ligands for the β_1_-adrenoceptor, thereby reducing the overall burden of side effects associated with the use of β-blockers.

## Abbreviations

CHO: Chinese hamster ovary
HBSS: Hanks’ balanced salt solution
[^3^H]-DHA: 1-[4,6-propyl-3H] dihydroalprenolol
NSB: non-specific binding.

## Author Contributions

Participated in research design: Sykes, Charlton, Veprintsev

Conducted experiments: Sykes, Reilly, Jiménez-Rosés

Performed data analysis: Sykes, Jiménez-Rosés, Reilly

Wrote or contributed to writing on manuscript: Sykes, Charlton, Jiménez-Rosés, Veprintsev, Reilly, Fairhurst

## Supplemental File

**Supplemental Figure 1.**
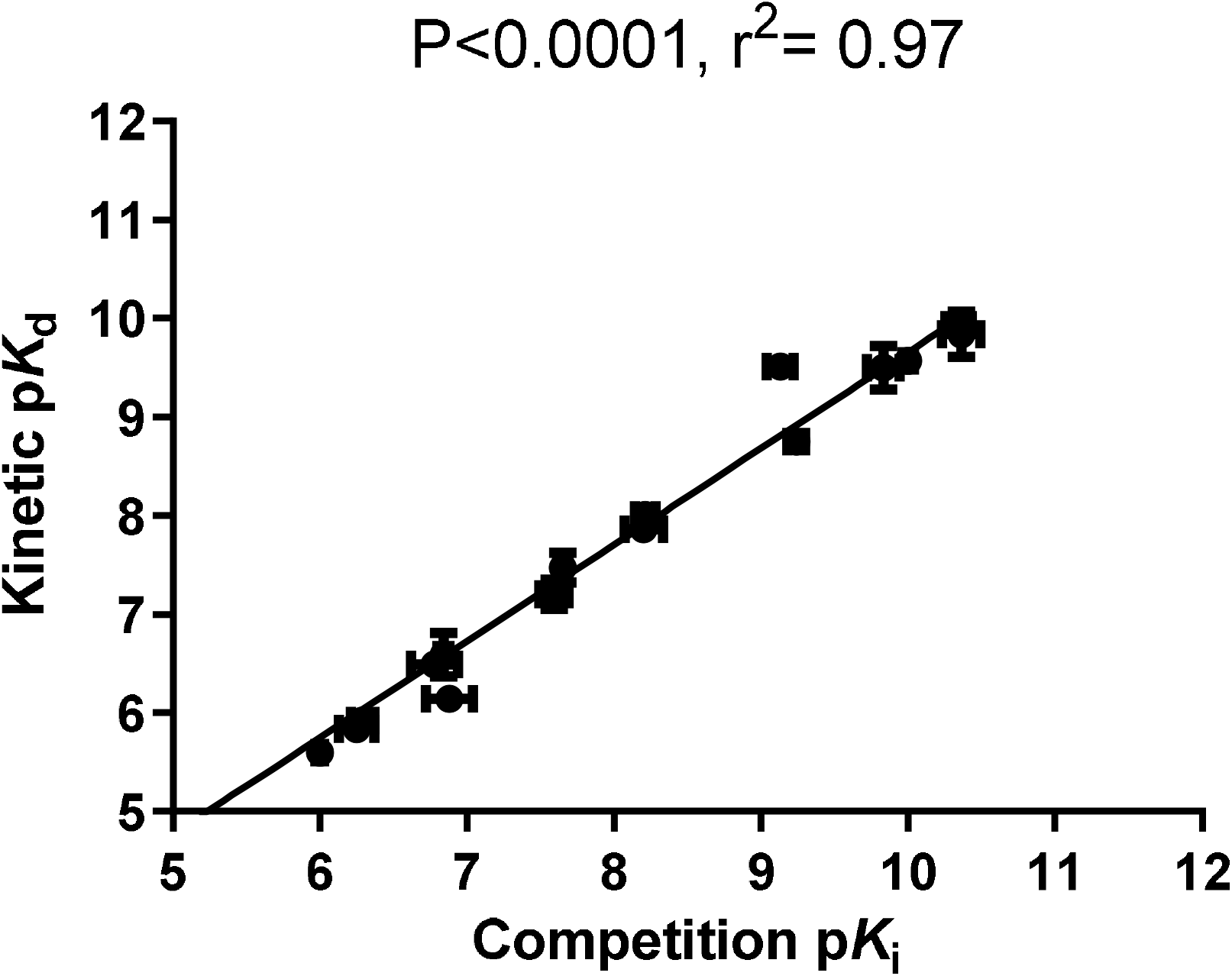
Correlating kinetically derived parameters of β_1_-adrenoceptor ligand. Correlation between p*K*_i_ and kinetically derived p*K*_i_ for the 17 test ligands. p*K*_i_ values were taken from [^3^H]-DHA competition binding experiments at equilibrium. The values composing the kinetically derived *K*_d_ (*k*_off_/*k*_on_) were taken from the experiments shown in Fig 4. All data used in these plots are detailed in Table 1. Data are presented as mean ± S.E.M. from three or more experiments.

**Supplemental Figure 2.**
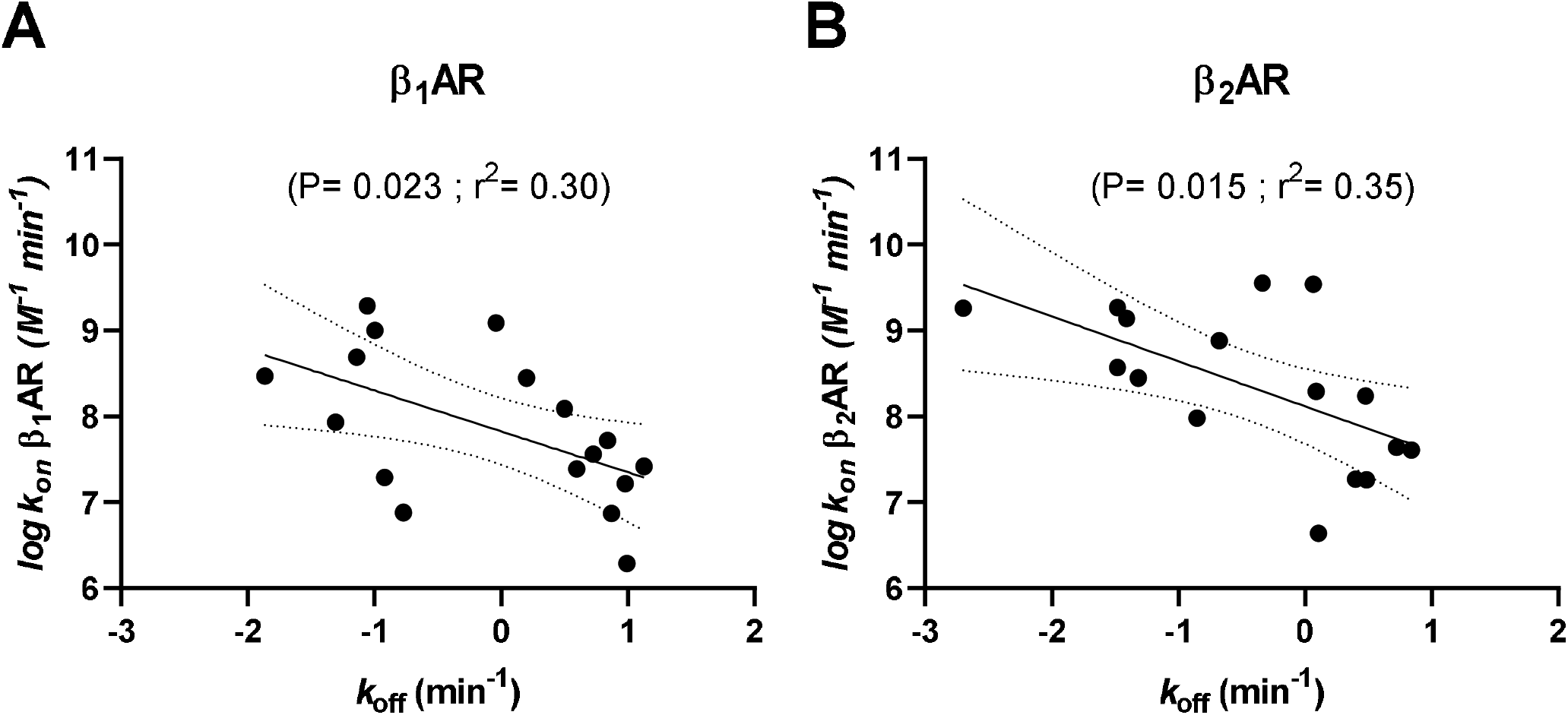
Relationship between *k*_off_ and *k*_on_ values for ligands targeting β-adrenoceptor subtypes 1 and 2. Kinetic values for ligands binding the β_2_ adrenoceptor were taken from Sykes et al., 2014.

